# Reverse Engineering Gene Networks Using Global-Local Shrinkage Rules

**DOI:** 10.1101/709741

**Authors:** Viral Panchal, Daniel Linder

## Abstract

Inferring gene regulatory networks from high-throughput ‘omics’ data has proven to be a computationally demanding task of critical importance. Frequently the classical methods breakdown due to the curse of dimensionality, and popular strategies to overcome this are typically based on regularized versions of the classical methods. However, these approaches rely on loss functions that may not be robust and usually do not allow for the incorporation of prior information in a straightforward way. Fully Bayesian methods are equipped to handle both of these shortcomings quite naturally, and they offer potential for improvements in network structure learning. We propose a Bayesian hierarchical model to reconstruct gene regulatory networks from time series gene expression data, such as those common in perturbation experiments of biological systems. The proposed methodology utilizes global-local shrinkage priors for posterior selection of regulatory edges and relaxes the common normal likelihood assumption in order to allow for heavy-tailed data, which was shown in several of the cited references to severely impact network inference. We provide a sufficient condition for posterior propriety and derive an efficient MCMC via Gibbs sampling in the Appendix. We describe a novel way to detect multiple scales based on the corresponding posterior quantities. Finally, we demonstrate the performance of our approach in a simulation study and compare it with existing methods on real data from a T-cell activation study.

## 1 Introduction

Methods for inferring gene networks attempt to probe the relationships between various cellular constituents like proteins, metabolites and gene products from data. These networks and pathways allow intracellular species to interact with each other, and external stimuli, so that the resulting coordinated molecular activity can support a wide range of cellular processes, like immune response [1], cell cycle, and death. Indeed, the importance of this underlying regulatory control structure is known to have implications in experimental and clinical biology.

The advances in high-throughput technologies, like the microarray and more recently next generation sequencing, has given rise to massive amounts of genomic data that can be used to learn these underlying regulatory networks [2, 3]. However, network inference has remained a significant challenge due to computational issues, detailed modeling assumptions, complicated likelihoods, and in general high-dimensionality [4].

A great deal of effort has been directed to developing methods for learning reaction network systems, which consists of a body of work that is too broad to fully review here, see [5]. Generally, most inference routines are either focused on structural inference or kinetic parameter estimation, with key modeling elements borrowed from a wide range of mathematical tools, like mutual information in [6], algebraic statistical models in [7, 8], ordinary differential equation (ODE) modeling of [9], and regularized objective functions [10], to name a few. From a statistical perspective, it is ideal to base inference on the likelihood function, since the Likelihood Principle has strong axiomatic foundations, and beyond that, estimates based on likelihoods exhibit optimal properties. The exact data likelihoods for stochastic reaction networks are usually intractable, since each reaction event is seldom recorded. While some work has been done using exact likelihoods, for instance in [11, 12, 13, 14], these methods are generally reserved for small systems of not more than a few molecular species types. Approximations to the exact processes, like the diffusion approximation [15, 16, 17] and the linear noise approximation (LNA) [18, 19, 20] trade some accuracy for computational speed. Both of these approximations become exact under limiting arguments [17], and in the case of LNA lead to a tractable Gaussian likelihood describing the process transition density.

The approximating stochastic differential equations (SDE)/ODEs provide a natural approach to gene network reconstruction through formulation of the inference problem via a regression type analysis. [21] developed a Bayesian reverse engineering strategy based on treating the data as an auto-regressive process, with point mass priors on the adjacency matrix indicating regulatory relationships. Although their data modeling assumption is frequently used for network inference, regressing total mRNA expression on time lagged expression values is problematic since transcription factors directly affect the rate of change of target mRNA and not the accumulated amount of target mRNA [6]. Recently, [22] cast the network reconstruction problem into a Bayesian regression framework as well. Unfortunately, their model regresses mRNA expression levels onto all other expression values at the same time point, making causal conclusions less straightforward. In addition, by treating the data as independent, statistical dependencies in transcriptional activity not due to the regulatory effects of measured quantities are not accounted for in their methodology. The method that we propose addresses both of these issues.

## 2 Methods

Our approach to inferring gene regulatory networks is based on a data model that utilizes a Euler-Maruyama discretization, see [23], to approximate the differentials of an SDE/ODE. This data modeling strategy allows the network inference based on SDEs and ODEs to proceed similar to a standard regression analysis, even though the inferred model is dynamical one. In combination with appropriate priors of a Bayesian model, the reformulation as a linear (statistical) model was shown to empirically produce good network estimates [4].

### 2.1 Expression data

Consider gene expression data, ***X***_*i*_(*t*_*j*_) ∈ ℝ^*d*^, denoting the expression values of *d* molecular species at time point *t_j_* for replicate *i*, where *i* = 1, …, *r* are independent experimental replications of the process measured at *t*_*j*_ *j* = 1, …, *m* time points, not necessarily equidistant. To form the response and design matrix, finite differences between time points are computed by

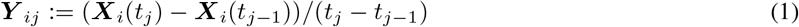

and then the response, or approximate differential ***Y***_*ij*_, can be regressed onto the lagged expression values ***X***_*i*_(*t*_*j*−1_). A natural way to perform network inference is to consider estimating the regulatory matrix of the following statistical model

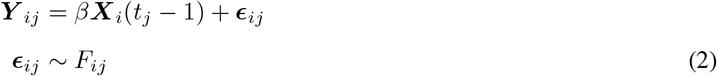

where *β* ∈ ℝ^*d×d*^ is the parameter of interest and *F*_*ij*_ is some *d*-dimensional probability measure. For large systems; i.e., when *d* is more than a few tens, some or many of the entries of *β* may be zero, this is often referred to as sparsity. Our Bayesian hierarchical model accounts for this by attempting to shrink the irrelevant entries in *β* to zero using priors we have specialized to the multivariate setting.

One pertinent observation about Eq. (2) is that it approximates a linear SDE/ODE. The proposed methodology easily extends to and is well defined in higher order systems, for instance those obtained by adding higher order terms like 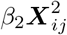 to the right-hand side of Eq. (2). This follows from a result we provide in the Appendix about a more general model, which will guarantee posterior propriety and an efficient Gibbs strategy, regardless of the order of the system. However, the nonlinear forms derived from single-cell biochemistry are not appropriate for data aggregated over cellular populations, like for instance in microarray and RNA-seq. The is because errors are incurred when commuting drift and expectation for nonlinear systems [4], so that averaging over populations will maintain consistency only in linear systems. Further, [4] report no observed improvements in network inference by including such extra basis functions over the linear model, although our approach does easily allow for such an extension. Further, they show that this linear model includes a number of existing approaches as special cases. They do note however that a crucial challenge is how to adequately account for heterogeneity due to uneven sampling intervals, which they show can lead to problems with inference. The commonly adopted strategy is to model these differences in variation by assuming that 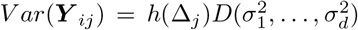, where *h*: ℝ^+^ → ℝ^+^ is a variance function and *D* is a diagonal matrix. However, the correct choice for such a variance function is impossible without knowledge of the unknown process. Our strategy attempts to learn this from the data.

### 2.2 Hierarchical model

To address these challenges, we propose a fully Bayesian procedure. Our proposed model takes the following form

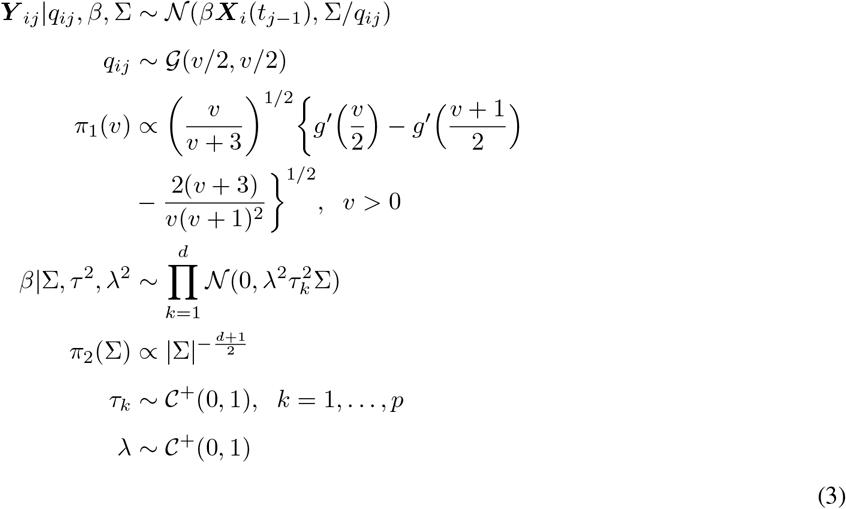

The motivation for choosing the terms in the above hierarchical model deserves some attention. We model the error terms, *ϵ*_*ij*_, as coming from the class of scale mixtures of multivariate normal distributions. We posit that this likelihood specification will be accurate in large volume and guards against potential heavy-tailed error. These have been observed in the various omics technologies, and we also show empirical evidence that our T-cell data exhibits this property. Independence and normality, conditional on the lagged expression and variance, follows from the Markov property and Gaussian transition density of the approximate process. The data likelihood model above will be a good approximation when the trajectories arise as observations from a stochastic reaction network in large volume or from a deterministic system, and then contaminated by heavy-tailed multivariate error. A key aspect of the formulation in Eq. (3) is the variance mixture component term, *q*_*ij*_, that can automatically adjust for uneven sampling intervals by mixing over scales of the normal distribution. Our choice of the Gamma prior on *q_ij_* leads to the multivariate Student’s-t likelihood for fixed degrees of freedom *v* [24]. We add another layer of hierarchy by assigning a non-informative prior on *v* to learn the tail heaviness from the data, instead of assigning it a fixed value a priori. We also assign a Jeffreys prior on the covariance matrix Σ. Our reliance on non-informative priors for these variance components of the likelihood are on purpose, since previous work has shown that inference is sensitive to these choices, and we prefer to learn these primarily from the data. Since the joint prior is then improper, checking for posterior propriety becomes imperative, and we give a sufficient condition in the Appendix for posterior propriety in a more general setting.

We assign independent scale mixture of multivariate normals to the columns of *β*. This is a multivariate extension of the class of global-local shrinkage priors, where 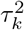 controls local shrinkage of the *k*^*th*^ column vector of *β* and *λ*^2^ controls global shrinkage across all parameters. The half-Cauchy distributions, 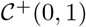, on the roots of the variance components produce marginal densities on *β* with infinites spikes at the origin and heavy tails that are able to separate signal from noise, see [25]. Importantly, although this prior is continuous, it has been shown to behave strikingly similar to the two-group model, or point mass mixture prior.

### 2.3 Posterior computation

An extremely useful and practical aspect of the hierarchical model in (3) is that it leads to an efficient computational strategy via data augmentation [26]. Here, we provide the Gibbs algorithm for posterior computation, and have made the corresponding R code implementation available online.

**Algorithm 1.**
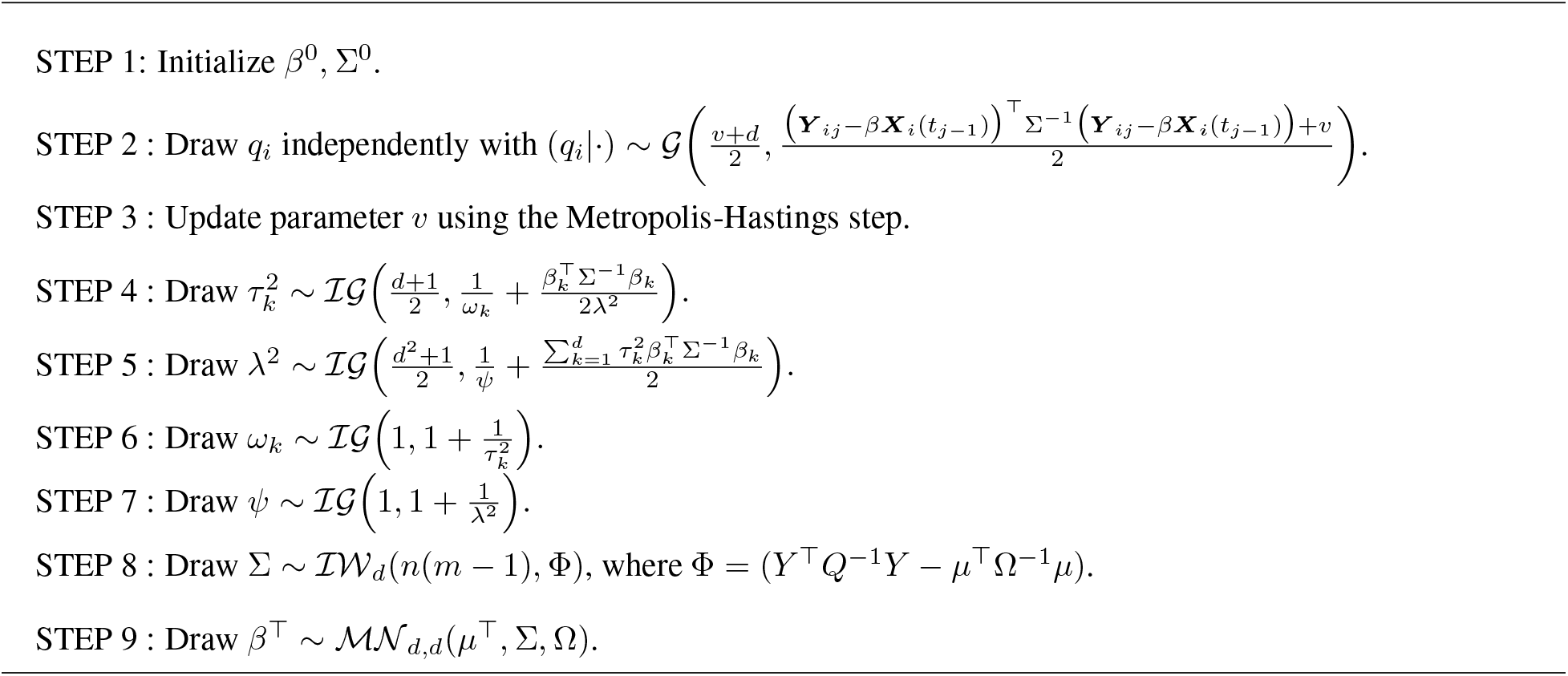
MCMC Algorithm

In our notation above, *Q* is the *r*(*m* − 1) × *r*(*m* − 1) diagonal matrix with elements *q*_*ij*_, *T* is the *d* × *d* diagonal matrix whose *k*^*th*^ element is 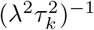, Ω = (***X***^T^*Q*^−1^***X*** + *T*^−1^)^−1^ and *μ* = Ω***X***^T^*Q*^−1^***Y***. The quantity ***X*** is the *r*(*m* − 1) × *p* design matrix with (*ij*)^*th*^ row ***X***_*i*_(*t*_*j*−1_) and ***Y*** is the *r*(*m* − 1) × *d* matrix whose (*ij*)^*th*^ row is ***Y***_*ij*_, for *i* = 1, …, *r* and *j* = 2, …, *m*. The term 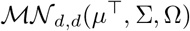 is the matrix normal distribution with location *μ* and scale matrices Σ and Ω. 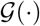 is a gamma distribution and 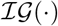 denotes inverse gamma distribution with respective parameters. 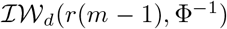 is the inverse-Wishart distribution with degrees of freedom *r*(*m* − 1) and scale matrix Φ^−1^.

### 2.4 Multiscaling

We outline here a novel method for using posterior quantities to both separate signal from noise, and also detect different scales in the signals. The idea behind multiscale clustering is to categorize parameters into clusters and use them to perform model selection subsequently. First, we calculate the mean and standard deviation for each coefficient from the posterior sample. Using the absolute value of the posterior mean of a coefficient, |*μ*_*β*_|, and |*μ*_*β*_|/*σ*_*β*_ we can form the 2-dimensional vector (|*μ*_*β*_|, |*μ*_*β*_|/*σ*_*β*_) to determine the different groups of coefficients. As an example, consider a reaction system with fast, slow, and superfluous regulatory interactions. These would correspond to large, small, and zero reaction rates/coefficients with three distinct clusters. Large |*μ*_*β*_|, and |*μ*_*β*_|/*σ*_*β*_ would represent fast reactions that are significant, while small |*μ*_*β*_| and large |*μ*_*β*_|/*σ*_*β*_ indicate slow but significant ones, the rest being either zero or not significant. If a researcher suspects more than two scales, additional numbers of clusters can be used for further scale detection. For instance, consider the absolute mean values of a hypothetical coefficient to be *β* = (1.2, 0.9, 1.5, 0.02, 0.012, 0.05, 0.01, 0.003, 4.5, 5.15, 4.2, 0.012, 0.033, 0.025). In this case, we have three obvious clusters: cluster 1 contains the values close to zero, cluster 2 contains (1.2, 0.9, 1.5), and cluster 3 with (4.5, 5.15, 4.2). We use K-means clustering to identify these different scales in subsequent analyses.

## 3 Simulation study

In this section, we design a simulation study to investigate how the proposed hierarchical model performs when the truth is known. We compare it to some standard statistical approaches across different metrics.

### 3.1 Parameter settings

We simulate time series gene expression profiles under the assumption of the following model:

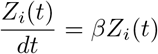

where *β* is the *d* × *d* regulatory matrix of the underlying true network. We set *d* = 25 so that *β* contains 25×25 = 625 unknown parameters of interest. For the reported results, we assume 10 true non-zero coefficients with values 0.91, 5.23, −1.04, 4.86, 4.5, 1.19, −0.81, −4.25, −4.23, 5.53 and set the rest to zero. This coincides with a fast scale of coefficients 5.23, 4.86, 4.5, −4.35. − 4.22, 5.53, a slower scale of coefficients 0.91, −1.04, 1.19, −0.81 and the zero ones. The true regulatory network structure is indicated in figure 3. Thus, the sparsity level here is 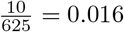. Each replicate has initial conditions randomly sampled from the *d* = 25 dimensional truncated normal distributed with mean 11 and variance 36, truncated below at 5 and above at 15. This is to simulate heterogeneity in the initial conditions between replications. We collect the ODE data at *m* = 6 randomly sampled time points between 0 and 5 and then contaminate them with noise; i.e., 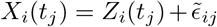, to arrive at the final data. We consider 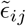 to be distributed as either normally distributed with variances (*σ*^2^) or Student’s t-distributed with degrees of freedom (*df*). The response, Y, is computed as discussed earlier by

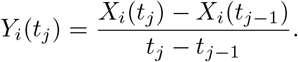

and the network inference is then based on estimating the coefficient/regulatory matrix of the multivariate regression model

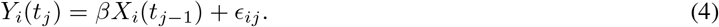

using ordinary least squares approach (OLS), Jeffreys’ non-informative prior (NIP), and our Bayesian hierarchical model (GLP). The OLS approach is perhaps the most frequently used classical method for estimation, but is known to have serious problems in sparse or data poor situations. The Jeffreys prior 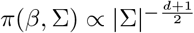 is a non-informative prior measure in our current setting, but it unfortunately leads to an improper posterior distribution under small replications. In the general multivariate regression setting, this happens when *n* < *p* + *d*, so that as the size of predictors *p* grows this is no longer an option. We prove in the Appendix that for our partially informative prior on *β* and Σ, a sufficient condition for propriety is *n* ≥ *d* in both the general setting and in our specialized setting.

**Figure. 1:**
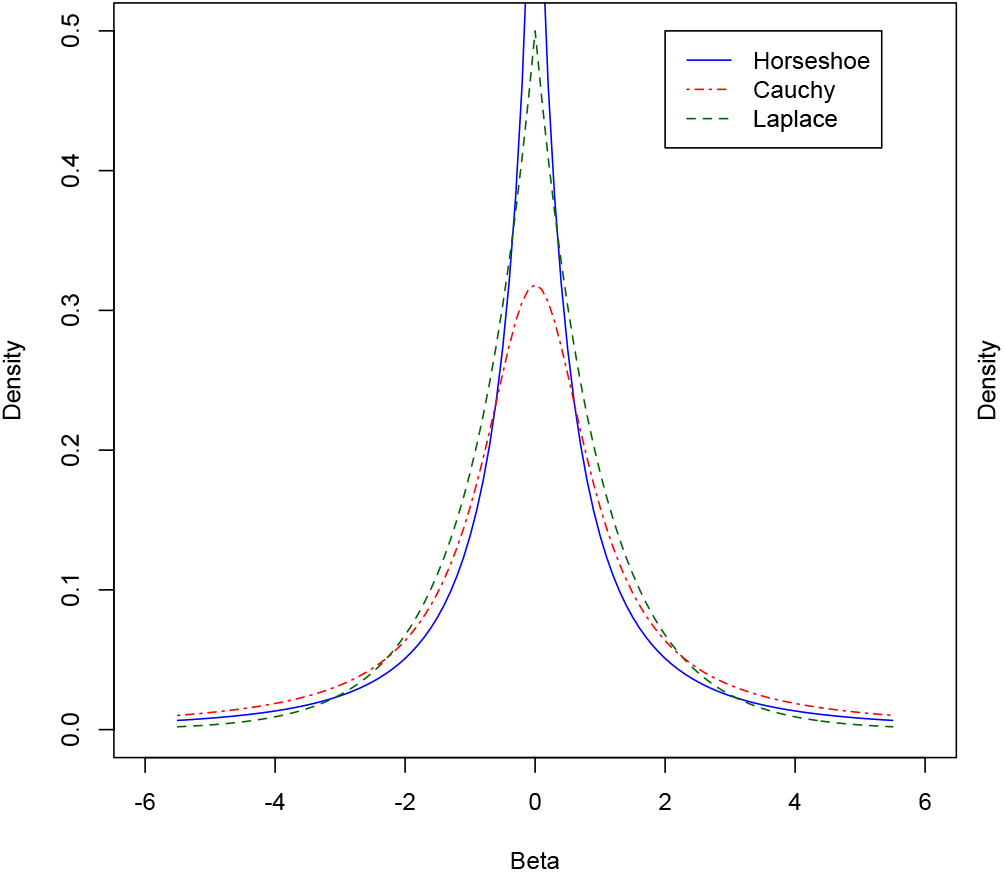
Comparison of prior densities

**Figure. 2:**
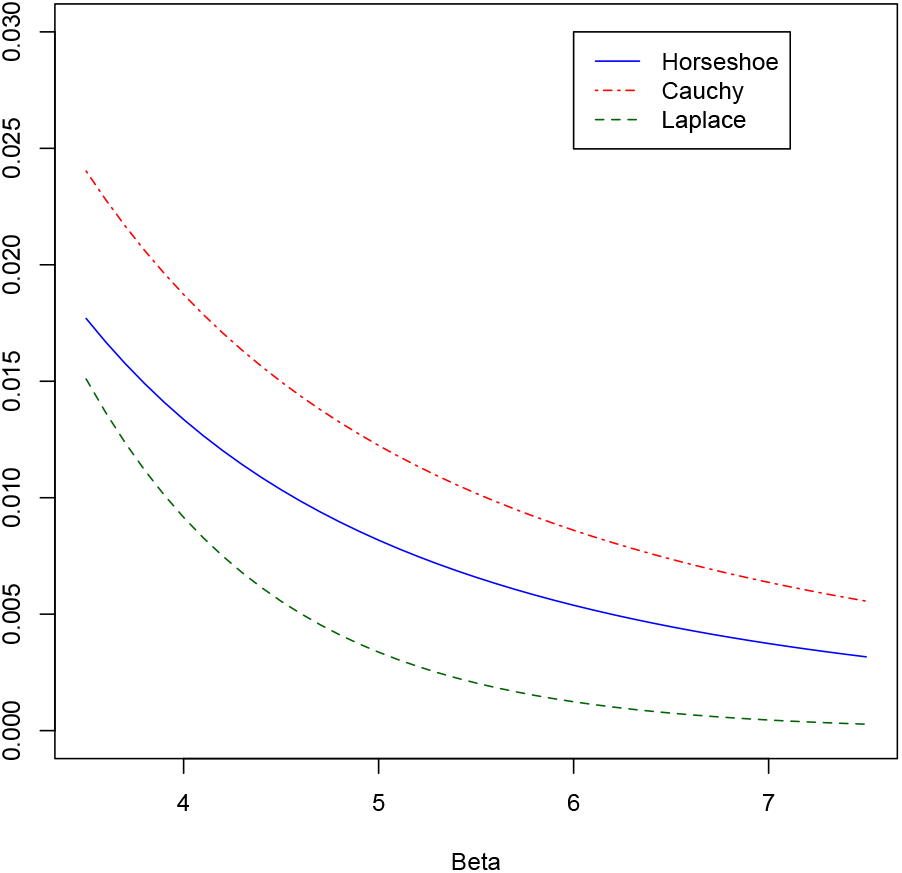
Tails of prior densities

**Figure. 3:**
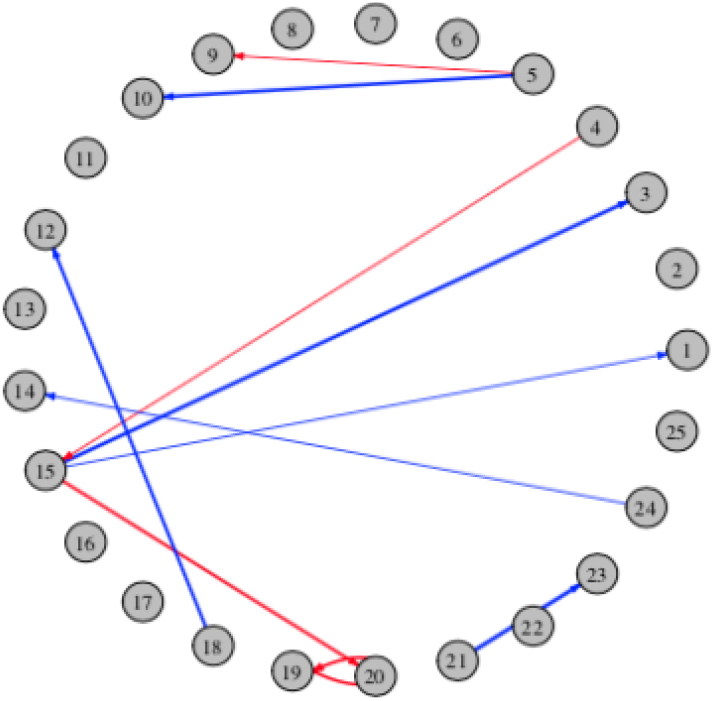
True network for simulation study with 25 nodes and 10 edges.

We note that the measurement time points are not equally spaced. The above error term, *ϵ*_*ij*_, is intended to capture the measurement errors 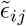 from the Euler data processing step, together with stochastic fluctuations that are heterogeneous across time, by accounting for this through the mixing of data likelihood over the error variance in the hierarchical model. Moreover, different combinations of *σ*^2^ and *df* are considered, as well differing numbers of replicates *r* to study low and high dimensional situations. The following combinations of *r*, *σ*^2^, and *df* are used in our study.

1. *r* ∈ {8, 12, 16} corresponding to *n* = *r* × (*m* − 1) ∈ {40, 60, 80}
2. *σ*^2^ ∈ {0.01, 0.1, 1}, for normal errors
3. *df* ∈ {3, 5, 10}, for student’s t errors

### 3.2 Simulation results

We compare the three approaches previously mentioned in terms of prediction accuracy, estimation and variable selection. For prediction accuracy, we calculate the prediction median absolute deviation (PMAD) between true and predicted responses. For parameter estimation, we compute the Frobenius norm (FN) between the true coefficient matrix and the estimated one. Table 1 summarizes both the estimation and prediction performance of all the methods under Student’s t error. In all examples, the global-local shrinkage prior significantly outperforms its competitors. We can see that when sample size decreases, FN and PMAD increase for all the methods, however, the GLP is less effected by drastic change in the sample size. It is also noted that with increasing *df*, FN and PMAD reduce in size for all methods as expected. We reason that the improved performance is due to the proposed model’s ability to handle both sparsity and heavier tailed, heterogeneous error. We report similar findings for normal errors (Table 3) with different measurement error variance *σ*^2^ = (0.01, 0.1, 1). The fully non-informative prior cannot be applied in the last case, because the posterior distribution is improper when *n* < *p* + *d*, note on our specialized setting since *p* = *d*, this implies impropriety when *n* < 50. We indicate this as NA in the subsequent tables. Additionally, we note that OLS gives erratic behavior because of identifiability issues, see Table 1.

**Table 1:** Simulation results for Student’s t error. The numbers are the averages of Frobenius norm (FN) and predicted median absolute deviation (PMAD) from 100 simulation replications for different scenarios of sample sizes and degrees of freedom.

| n | df | FN |  |  | PMAD |  |  |
| --- | --- | --- | --- | --- | --- | --- | --- |
|  |  | GLP | OLS | NIP | GLP | OLS | NIP |
| 80 | 3 | 5.37 | 13.88 | 13.19 | 1.85 | 3.06 | 2.90 |
|  | 5 | 4.43 | 12.74 | 12.45 | 1.62 | 2.69 | 2.58 |
|  | 10 | 4.08 | 12.45 | 12.52 | 1.50 | 2.53 | 2.44 |
| 60 | 3 | 5.79 | 17.35 | 17.26 | 1.89 | 3.74 | 3.73 |
|  | 5 | 4.75 | 16.12 | 16.48 | 1.67 | 3.42 | 3.40 |
|  | 10 | 4.44 | 16.10 | 16.51 | 1.54 | 3.22 | 3.21 |
| 40 | 3 | 7.56 | 24.83 | NA | 2.01 | 5.37 | NA |
|  | 5 | 6.71 | 25.79 | NA | 1.79 | 5.10 | NA |
|  | 10 | 6.24 | 24.59 | NA | 1.63 | 4.76 | NA |

To illustrate the effect of shrinking coefficients using the global-local shrinkage prior in the proposed variable selection procedure in 22.4, we plot the estimated stable edges over 100 iterations in figure 4. These are the edges appearing in at least half of the iterations. We also select the stable edges in the real data analysis in this way; i.e., as those edges from the full data analysis that were also present in at least half of the inferred networks from the bootstrap resamples, see below. From this figure, we can see that as sample size increases our method tends to identify a higher number of true edges than other methods. Although we see some falsely identified edges, they are comparatively less than other methods. The method based on the non-informative prior performs worse than the proposed method, but better than the OLS in terms of network identification. We quantify the network inference capabilities of different methods by reporting the average true positive rate (TPR) as the number of edges correctly identified by the method, the false discovery rate (FDR) as the ratio of falsely identified edges and total number of selected edges, and the false positive rate (FPR) as the number of incorrectly selected edges, as illustrated in tables 2 and 4. The formulas for each of these evaluation criteria are given as

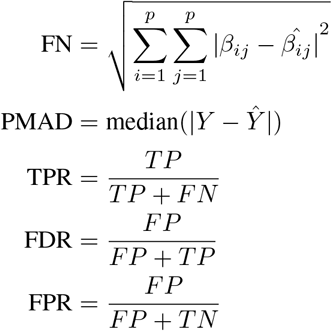

**Table 2:** Simulation results for student’s t error. The numbers are the averages of true positive rate (TPR), false discovery rate (FDR), and false positive rate (FPR) from 100 simulation replications for different scenarios of sample sizes and degrees of freedom.

| n | df | TPR |  |  | FDR |  |  | FPR |  |  |
| --- | --- | --- | --- | --- | --- | --- | --- | --- | --- | --- |
|  |  | GLP | OLS | NIP | GLP | OLS | NIP | GLP | OLS | NIP |
| 80 | 3 | 0.950 | 0.924 | 0.940 | 0.554 | 0.894 | 0.881 | 0.021 | 0.131 | 0.117 |
|  | 5 | 0.959 | 0.954 | 0.961 | 0.441 | 0.875 | 0.866 | 0.013 | 0.114 | 0.106 |
|  | 10 | 0.978 | 0.947 | 0.954 | 0.405 | 0.868 | 0.864 | 0.012 | 0.106 | 0.102 |
| 60 | 3 | 0.915 | 0.870 | 0.882 | 0.587 | 0.919 | 0.914 | 0.024 | 0.168 | 0.161 |
|  | 5 | 0.929 | 0.871 | 0.883 | 0.516 | 0.911 | 0.910 | 0.017 | 0.153 | 0.153 |
|  | 10 | 0.948 | 0.891 | 0.888 | 0.486 | 0.902 | 0.905 | 0.016 | 0.143 | 0.147 |
| 40 | 3 | 0.818 | 0.785 | NA | 0.622 | 0.949 | NA | 0.025 | 0.247 | NA |
|  | 5 | 0.856 | 0.786 | NA | 0.610 | 0.947 | NA | 0.024 | 0.241 | NA |
|  | 10 | 0.853 | 0.811 | NA | 0.566 | 0.947 | NA | 0.020 | 0.248 | NA |

**Table 3:** Simulation results for normal error. The numbers are the averages of true positive rate (TPR), false discovery rate (FDR), and false positive rate (FPR) from 100 simulation replications for different scenarios of sample sizes and variances.

| n | $\sigma^2$ | TPR | | | FDR | | | FPR | | |
| --- | --- | --- | --- | --- | --- | --- | --- | --- | --- | --- |
|  |  | GLP | OLS | NIP | GLP | OLS | NIP | GLP | OLS | NIP |
| 80 | 0.01 | 1.000 | 0.953 | 0.930 | 0.282 | 0.804 | 0.801 | 0.006 | 0.066 | 0.064 |
|  | 0.10 | 1.000 | 0.981 | 0.978 | 0.284 | 0.795 | 0.790 | 0.006 | 0.068 | 0.065 |
|  | 1.00 | 0.983 | 0.968 | 0.963 | 0.370 | 0.854 | 0.846 | 0.010 | 0.097 | 0.091 |
| 60 | 0.01 | 1.000 | 0.760 | 0.739 | 0.278 | 0.889 | 0.884 | 0.006 | 0.104 | 0.098 |
|  | 0.10 | 0.999 | 0.884 | 0.883 | 0.286 | 0.887 | 0.885 | 0.007 | 0.119 | 0.117 |
|  | 1.00 | 0.948 | 0.872 | 0.869 | 0.449 | 0.894 | 0.891 | 0.014 | 0.125 | 0.119 |
| 40 | 0.01 | 0.998 | 0.459 | NA | 0.290 | 0.921 | NA | 0.007 | 0.088 | NA |
|  | 0.10 | 0.980 | 0.784 | NA | 0.327 | 0.935 | NA | 0.008 | 0.189 | NA |
|  | 1.00 | 0.875 | 0.807 | NA | 0.564 | 0.946 | NA | 0.020 | 0.240 | NA |

**Table 4:** Simulation results for normal error. The numbers are the averages of Frobenius norm (FN) and predicted median absolute deviation (PMAD) from 100 simulation replications for different scenarios of sample sizes and variances.

| n | $\sigma^2$ | FN | | | PMAD | | |
| --- | --- | --- | --- | --- | --- | --- | --- |
|  |  | GLP | OLS | NIP | GLP | OLS | NIP |
| 80 | 0.01 | 3.55 | 25.89 | 28.09 | 0.15 | 1.29 | 1.32 |
|  | 0.10 | 3.57 | 13.56 | 13.65 | 0.43 | 1.48 | 1.45 |
|  | 1.00 | 4.04 | 12.03 | 12.24 | 1.37 | 2.38 | 2.30 |
| 60 | 0.01 | 3.73 | 58.22 | 63.70 | 0.16 | 2.85 | 2.89 |
|  | 0.10 | 3.77 | 22.73 | 23.65 | 0.45 | 2.77 | 2.76 |
|  | 1.00 | 4.24 | 16.04 | 16.26 | 1.42 | 3.13 | 3.12 |
| 40 | 0.01 | 4.28 | 119.28 | NA | 0.17 | 4.68 | NA |
|  | 0.10 | 4.29 | 45.08 | NA | 0.49 | 4.53 | NA |
|  | 1.00 | 5.35 | 24.52 | NA | 1.51 | 4.73 | NA |

**Figure. 4:**
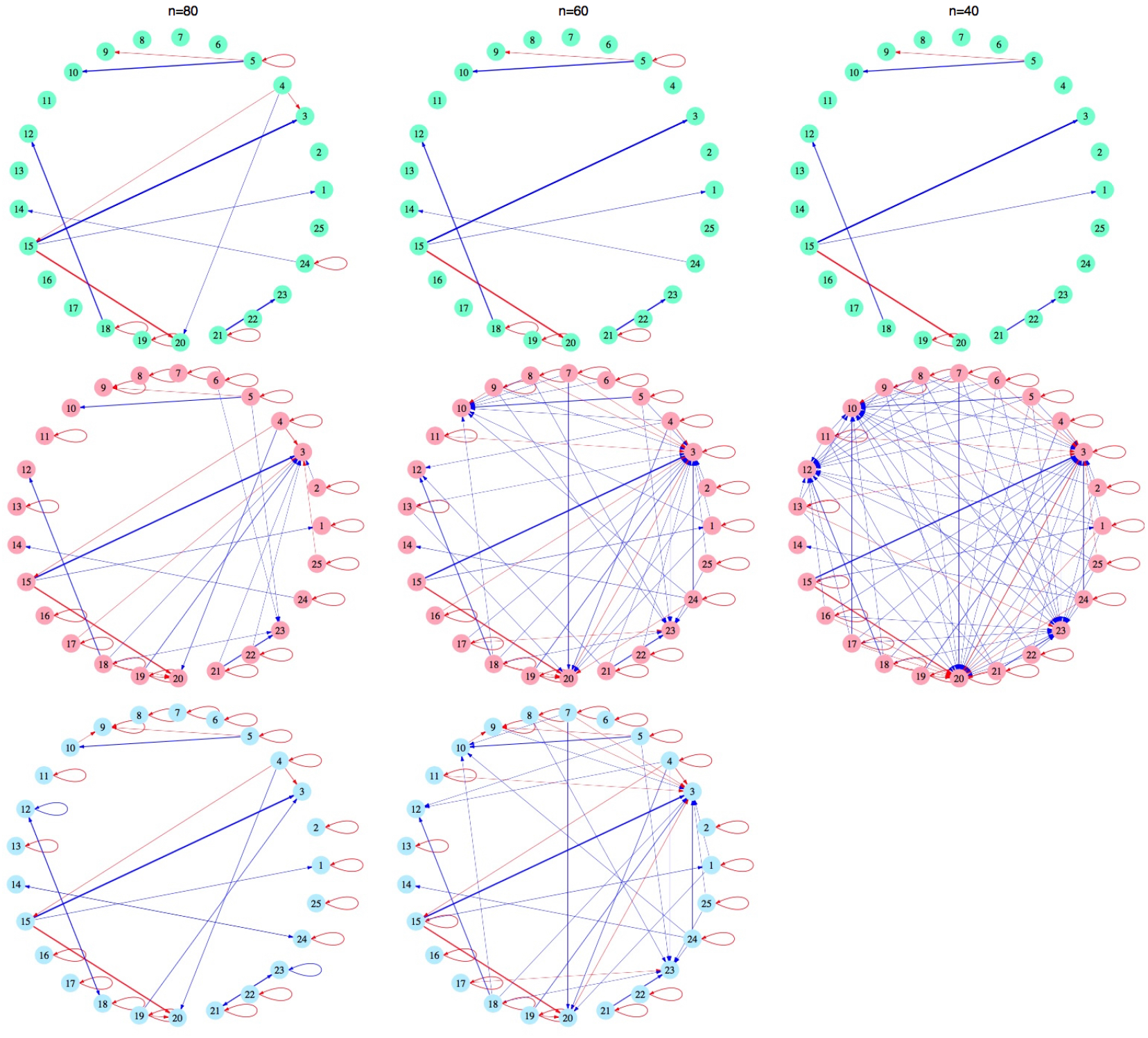
Inferred networks for simulation example with global-local prior (GLP) (green), ordinary least squares (OLS) method (pink) and non-informative prior (NIP) (blue). Plots were generated using the R CRAN package igraph.

Inspection of tables 2 and 4 reveals that GLP has markedly improved edge selection with the highest TPR, and least FPR and FDR over the other methods and in all the situations considered. These findings illustrate that not only is inference under the GLP hierarchy valid, but that it is robust across a range of sample sizes. However, when *n* approaches *d*, the posterior propriety threshold, GLP based inference becomes worse across the range of metrics, as expected. We again remark that inference via GLP is valid as long as *n* ≥ *d*. In summary, these simulation results show that the proposed method performs consistently better than other methods, and is robust to sample size and measurement error.

## 4 T-cell activation data - Reverse engineering gene regulatory network

In this section, we use the proposed methodology to infer a gene regulatory network from the time series microarray data for T-cell activation presented in [27]. The data originates from an experiment performed to identify the response of human T-cell line to PMA and ionomycin treatment. The authors investigated the expression of 88 genes using complementary DNA (cDNA) across 10 time points. Human T-cell line cells were treated with PMA and ionomycin, and cells were collected at the following time points after treatment: 0, 2, 4, 6, 8, 18, 24, 32, 48, and 72 hours. Then, Fluorescence-Activated Cell Scanning (FACS) analysis was performed to measure T-cell expression and activation markers.

Two identical experiments were conducted on two sets of microarrays. The first experiment consisted of microarrays representing 34 replications, with the second experiment containing 10 replications. The authors pre-selected 58 genes out of 88 genes after filtering out gene expression values based on a pre-defined threshold value. In addition, normalization and log-transformation were performed to minimize systematic variation due to experimental situations and to normalize the data.

We consider this pre-processed data on all 44 replications and 58 genes for the final analysis. The design matrix, ***X***, and the generated response matrix, ***Y***, are computed according to Eq. (1). We perform preliminary diagnostics for the error terms by calculating residuals from a least squares fit to the data; i.e., generated response regressed onto the design matrix. We provide some of the corresponding Q-Q plots in figure 5 for a subset of four different expression types. The figure indicates that the marginal error distributions are nearly symmetric, but exhibit very heavy tails, and indeed, the formal statistical tests of multivariate normality all fail (p-values < 0.0001).

**Figure. 5:**
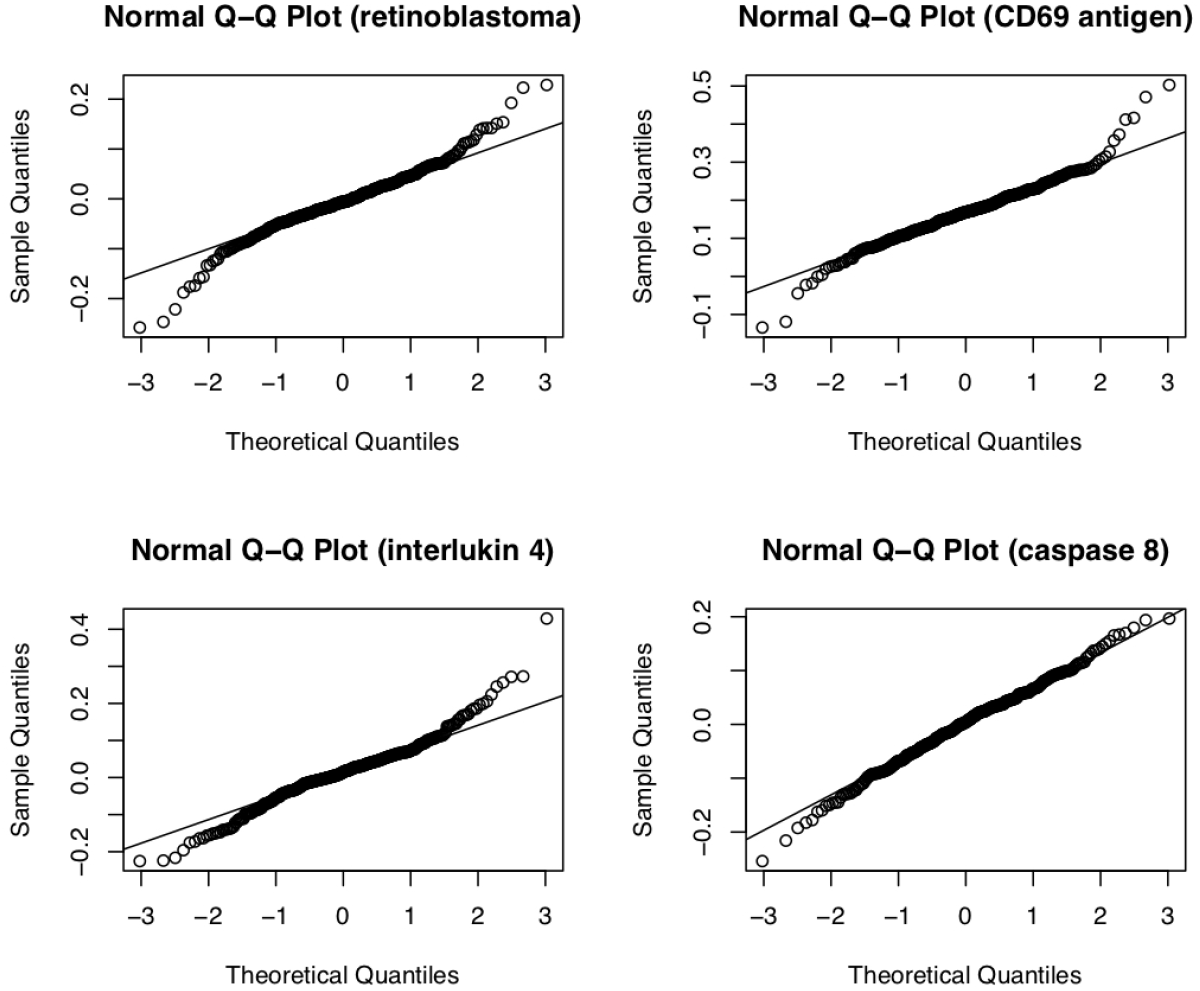
Q-Q plots

The final gene regulatory network that we report is computed by first fitting our Bayesian hierarchical model to the full data, and then selecting from the inferred edges the stable edges; i.e., those edges that showed up in at least 50% of 100 bootstrap resamples, see figure 6. The multiscaling strategy based on the posterior were used in each of the 100 resamples to compute the network for that sample. The R package ‘igraph’ was used to plot the inferred gene regulatory network in figure 6. Next we discuss some of the biological validity of these findings.

**Figure. 6:**
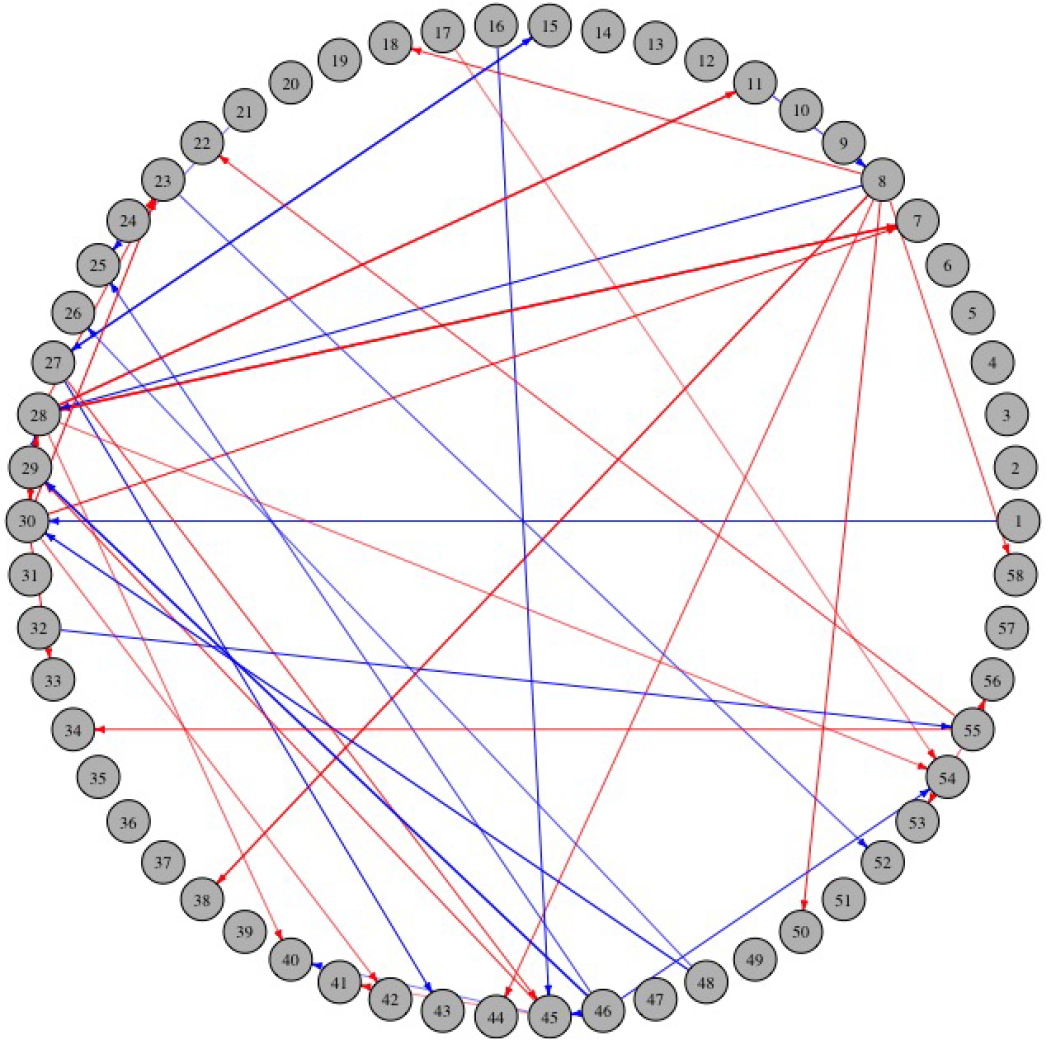
Directed gene regulatory network representing the elements of the regulatory matrix.

The process of T-cell activation and response comprises of four main steps: adhesion, inflammation, differentiation, and apoptosis. The network in figure 6 consists of genes involved in all four of these steps. In the inferred network, ZNFN1A1 (gene 8), EGR1 (gene 28), MCL1 (gene 30), FYB (gene 45), IL2RG (gene 46), and IL3RA (gene 55) have the highest number of connections. ZNFN1A1 encodes a zinc finger protein called Ikaros, which is essential for T-cell proliferation and differentiation [28]. ZNFN1A1 inhibits cell death by negatively influencing the expression of CASP4 and CASP8. Also, JUND affects T-cell proliferation by upregulating ZNFN1A1. We identified three FYB-regulated genes that were also found in Rangel et al, which are involved in activation (GATA3), proliferation (API2), and inflammation (IL2RG). The FYB gene plays a vital role in T-cell proliferation by activating the API2 gene, which functions in the cell by inhibiting caspases in the induction of cell death. EGR-1 is an important gene in the regulation of cell growth, differentiation, and apoptosis. Our model demonstrates an interesting association between Early Growth Response gene (EGR1) with the API1 gene. EGR1 was determined as an early response gene influenced by mitogenic activation [29, 27]. According to our model, EGR1 negatively regulates API2 which is a gene responsible for inhibiting cell apoptosis. Furthermore, EGR1 inhibits cell growth and differentiation by suppressing genes CD69, JUND, and JUNB. Several studies have reported the functional role of EGR-1/JUN complex in cell apoptosis [30]. Other examples of genes that appear linked in the inferred network include IL2RG, IL3RA, CD69, TRAF5, and IL16, which are involved in the production of cytokines and regulate the inflammatory response [31].

## 5 Concluding remarks

In this work, we have proposed a Bayesian hierarchical model using multivariate global-local shrinkage priors to infer gene networks. The proposed method handles heavy-tailed data by assuming a multivariate heavy-tailed data likelihood that mixes over Gaussian variance components. Our simulation study demonstrates superior performance of the proposed method in terms of edge selection, coefficient estimation, and outcome prediction as compared to other approaches and in a variety of different situations. Our real data analysis reveals edges that have been confirmed experimentally and suggests interesting novel relationships for T-cell activation.

We have shown that the proposed method performs well in the high-dimensional situation, and justify its use in this case by providing sufficient conditions for posterior propriety. This is essential to check when the joint prior distribution is not proper since it is required for valid Bayesian inference. When *n* ≥ *d*, the proposed hierarchical model has a proper posterior, and importantly this condition makes it straightforward to extend our model to arbitrary system types through simple basis expansion. In contrast, the fully non-informative prior yields an improper posterior distribution in high dimensions. Another contribution of our work is the multivariate extension of the global-local shrinkage prior. We are not aware of any previous work that integrates global-local shrinkage rules into the multivariate setting to reverse engineer gene networks. In the biological context, it is often assumed that many biological networks, such as the one under consideration, are indeed sparse. Thus, in the case of true underlying sparsity, the additional information encoded by the global-local prior significantly improves inference. In this paper, we have demonstrated in the simulations and real data application that the use of global-local shrinkage prior indeed produces such sparsity *a posteriori*.

Bayesian approaches often have computational challenges due to the required MCMC sampling in complicated or high dimensional models, but a new interpretation of the horseshoe prior [32] has made our posterior sampling procedure much more straightforward and computationally efficient. Although we alleviate a good deal of the MCMC tuning issues by deriving conjugate full conditional distributions for all parameters, naturally the sampling can become slow with increasing model size. We are currently exploring strategies to gain even more computational efficiency by considering alternative representations of certain full conditionals in the sampler.

## Acknowledgements

The authors would like to thank Bani K. Mallick for helpful discussions about the multiscale clustering strategy.

## Data Accessibility

The data used in this study is publicly available in the open source software R.

