## Supplemental Material for "Reverse Engineering Gene Networks Using Global-Local Shrinkage Rules"

### Supplementary Material

Viral Panchal and Daniel F. Linder

Medical College of Georgia, Augusta University

#### 1 Proof of posterior propriety

In this section, we give conditions for posterior propriety of $(\beta, \Sigma)$ under the full hierarchical model as shown below.

$$\begin{aligned}
Y_{ij}|q_{ij}, \beta, \Sigma &\sim \mathcal{N}(\beta X_i(t_{j-1}), \Sigma/q_{ij}) \\
q_{ij} &\sim \mathcal{G}(v/2, v/2) \\
\pi_1(v) &\propto \left(\frac{v}{v+3}\right)^{1/2} \left\{ g'\Big(\frac{v}{2}\Big) - g'\Big(\frac{v+1}{2}\Big) - \frac{2(v+3)}{v(v+1)^2} \right\}^{1/2}, \;\; v > 0 \\
\beta|\Sigma, \tau^2, \lambda^2 &\sim \prod_{k=1}^{p} \mathcal{N}(0, \lambda^2 \tau_k^2 \Sigma) \\
\pi_2(\Sigma) &\propto |\Sigma|^{-\frac{d+1}{2}} \\
\tau_k &\sim \mathcal{C}^+(0, 1), \;\; k = 1, \ldots, p \\
\lambda &\sim \mathcal{C}^+(0, 1).
\end{aligned} \tag{1}$$

Let us denote

$$c(y) = \int_{\mathbb{R}_+^{p+2}} \int_S \int_{\mathbb{R}^{dp}} f(y|\beta, \Sigma)\pi(\beta|\Sigma, \tau, \lambda)\pi(\Sigma)\pi(\tau)\pi(\lambda)\pi(v) \;\; d\beta d\Sigma d\tau d\lambda dv, \tag{2}$$

where $S$ represents the space of $d \times d$ positive definite matrices. To preserve notational simplicity, we use $\pi(\cdot)$ to denote prior densities, although it should be understood that each density corresponds to the appropriate term in the hierarchy (1). For posterior propriety, it is necessary that $c(y) < \infty$. When $c(y)$ is finite, the posterior density of $(\beta, \Sigma)$ is written as

$$\pi(\beta, \Sigma, \tau, \lambda, v|y) = \frac{f(y|\beta, \Sigma)\pi(\beta|\Sigma, \tau, \lambda)\pi(\Sigma)\pi(\tau)\pi(\lambda)\pi(v)}{c(y)}. \tag{3}$$

Our proof of posterior propriety is based on adaptations to the one given in [1] to account for the global-local scale mixture of normal priors on regression coefficients instead of the flat prior given there. We emphasize that the fully non-informative prior considered in [1] is not suitable in a high dimensional setting since it leads to improper posteriors when $n < d + p$, which is exactly the high-dimensional case under consideration, i.e. $p > n$.

**Proposition 1.** *Denote $\mu^{dn}$ as Lebesgue measure on $\mathbb{R}^{dn}$. If $n \geq d$, then $c(y) < \infty \quad a.e.(\mu^{dn})$, where a.e. is almost everywhere.*

*Proof.* Consider the missing data model consisting of regression data $y$, and the missing observations from the mixing density $h$. Given $(\beta, \Sigma)$, let $\{y_i, q_i\}, i = 1, ..., n$ be the independent and identically distributed pairs such that

$$\begin{aligned} y_i|q_i, \beta, \Sigma &\sim N_d(\beta^\top x_i, \Sigma/q_i) \\ q_i &\sim h. \end{aligned} \tag{4}$$

The joint density of $\{y, q\}$ can be expressed as

$$f(y, q|\beta, \Sigma) = \prod_{i=1}^{n} f(y_i|q_i, \beta, \Sigma)h(q_i), \tag{5}$$

and the marginal density of $y$ can be written as

$$\begin{aligned} f(y|\beta, \Sigma) &= \int_{\mathbb{R}_+^n} f(y, q|\beta, \Sigma)\, dq \\ &= \int_{\mathbb{R}_+^n} \prod_{i=1}^{n} f(y_i|q_i, \beta, \Sigma)h(q_i)\, dq_i \\ &= \prod_{i=1}^{n} \int_0^\infty \frac{(q_i)^{\frac{d}{2}}}{(2\pi)^{\frac{d}{2}}|\Sigma|^{\frac{1}{2}}} \exp\Big\{ -\frac{q_i}{2}(y_i - \beta^\top x_i)^\top \Sigma^{-1}(y_i - \beta^\top x_i)\Big\} h(q_i)\, dq_i. \end{aligned} \tag{6}$$

To obtain $c(y)$, we integrate out $\beta, \Sigma$, $q$, $\tau$, and $\lambda$ using the specified distributions in the hierarchy (1). From equation (2) and applying Fubini's theorem, we can express $c(y)$ as

$$\begin{aligned} c(y) = \int_{\mathbb{R}_+^{n+p+2}} \int_S \int_{\mathbb{R}^{dp}} & f(y, q|\beta, \Sigma)\pi(\beta|\Sigma, \tau, \lambda)\pi(\Sigma)\pi(\tau)\pi(\lambda)\pi(v) \\ & \times\, d\beta\, d\Sigma\, dq\, d\tau\, d\lambda\, dv. \end{aligned} \tag{7}$$

First, we simplify two of the seven integrals on the right-hand side of equation (7) as follows:

$$\begin{aligned} & f(y,q|\beta,\Sigma)\pi(\beta|\Sigma,\tau,\lambda)\pi(\Sigma) \\ = & \frac{1}{(2\pi)^{\frac{nd}{2}}|\Sigma|^{\frac{n}{2}}}\left[\prod_{i=1}^{n} q_i^{\frac{d}{2}}\exp\left\{-\frac{q_i}{2}(\beta^\top x_i - y_i)^\top\Sigma^{-1}(\beta^\top x_i - y_i)\right\}\right]\times \\ & \frac{1}{(2\pi)^{\frac{pd}{2}}|\Sigma|^{\frac{p}{2}}}\left[\prod_{k=1}^{p}(\lambda^2\tau_k^2)^{-\frac{d}{2}}\exp\left\{-\frac{(\lambda^2\tau_k^2)^{-1}}{2}\beta_k^\top\Sigma^{-1}\beta_k\right\}\right]\left[\prod_{i=1}^{n}h(q_i)\right]|\Sigma|^{-\frac{d+1}{2}} \\ = & \frac{|Q|^{-\frac{d}{2}}|T|^{-\frac{d}{2}}}{(2\pi)^{\frac{nd+pd}{2}}|\Sigma|^{\frac{n+p+d+1}{2}}}\left[\exp\left\{-\frac{1}{2}\sum_{i=1}^{n}q_i(\beta^\top x_i - y_i)^\top\Sigma^{-1}(\beta^\top x_i - y_i)\right\}\right]\times \\ & \left[\exp\left\{-\frac{1}{2}\sum_{k=1}^{p}\beta_k^\top(\lambda^2\tau_k^2\Sigma)^{-1}\beta_k\right\}\right]\left[\prod_{i=1}^{n}h(q_i)\right], \end{aligned} \tag{8}$$

where $\beta$ is the $p \times d$ coefficient matrix whose $k^{th}$ row is $\beta_j^\top$, $Q$ represents the $n \times n$ diagonal matrix whose $i^{th}$ diagonal entry is $q_i^{-1}$, and $T$ denotes the $p \times p$ diagonal matrix whose $k^{th}$ diagonal element is $(\lambda^2\tau_k^2)$. Now considering -2$\times$ the exponent term of equation (8) gives

$$\begin{aligned} & \sum_{i=1}^{n}q_i(\beta^\top x_i - y_i)^\top\Sigma^{-1}(\beta^\top x_i - y_i) + \sum_{k=1}^{p}(\lambda^2\tau_k^2)^{-1}\beta_k^\top\Sigma^{-1}\beta_k \\ = & \sum_{i=1}^{n}q_i\mathrm{tr}\Big[(\beta^\top x_i - y_i)^\top\Sigma^{-1}(\beta^\top x_i - y_i)\Big] + \sum_{k=1}^{p}(\lambda^2\tau_k^2)^{-1}\mathrm{tr}\Big[\beta_k^\top\Sigma^{-1}\beta_k\Big] \\ = & \mathrm{tr}\Big[\Sigma^{-1}(\beta^\top - \mu^\top)\Omega^{-1}(\beta^\top - \mu^\top)^\top\Big] + \mathrm{tr}\Big[\Sigma^{-1}(Y^\top Q^{-1}Y - \mu^\top\Omega^{-1}\mu)\Big], \end{aligned} \tag{9}$$

where $\Omega = (X^\top Q^{-1}X + T^{-1})^{-1}$ and $\mu = (X^\top Q^{-1}X + T^{-1})^{-1}X^\top Q^{-1}Y$. So,

$$f(y,q|\beta,\Sigma)\pi(\beta|\Sigma,\tau,\lambda)\pi(\Sigma) \propto \exp\left\{-\frac{1}{2}\mathrm{tr}\Big[\Sigma^{-1}(\beta^\top - \mu^\top)\Omega^{-1}(\beta^\top - \mu^\top)^\top\Big]\right\}. \tag{10}$$

From the density of the matrix normal distribution,

$$\int_{\mathbb{R}^{dp}}\exp\left\{-\frac{1}{2}\mathrm{tr}\Big[\Sigma^{-1}(\beta^\top - \mu^\top)\Omega^{-1}(\beta^\top - \mu^\top)^\top\Big]\right\}d\beta^\top = (2\pi)^{\frac{pd}{2}}|\Sigma|^{\frac{p}{2}}|\Omega|^{\frac{d}{2}}. \tag{11}$$

Thus, after marginalizing over $\beta$, Equation (8) becomes

$$\begin{aligned} & \int_{\mathbb{R}^{dp}} f(y,q|\beta,\Sigma)\pi(\beta|\Sigma,\tau,\lambda)\pi(\Sigma)\,d\beta^\top \\ = & \frac{|Q|^{-\frac{d}{2}}|T|^{-\frac{d}{2}}|\Omega|^{\frac{d}{2}}}{(2\pi)^{\frac{nd}{2}}|\Sigma|^{\frac{n+d+1}{2}}}\exp\left\{-\frac{1}{2}\mathrm{tr}\Big[\Sigma^{-1}(Y^\top Q^{-1}Y - \mu^\top\Omega^{-1}\mu)\Big]\right\}\left[\prod_{i=1}^{n}h(q_i)\right]. \end{aligned} \tag{12}$$

That the matrix $Y^\top Q^{-1}Y - \mu^\top \Omega^{-1}\mu$ is positive definite $a.e.(\mu^{dn})$ follows from expressing $Y^\top Q^{-1}Y - \mu^\top \Omega^{-1}\mu$ as

$$\begin{aligned} &Y^\top Q^{-1}Y - \mu^\top \Omega^{-1}\mu \\ &= \; Y^\top Q^{-\frac{1}{2}}\Big[I - Q^{-\frac{1}{2}}X(X^\top Q^{-1}X + T^{-1})^{-1}X^\top Q^{-\frac{1}{2}}\Big]Q^{-\frac{1}{2}}Y. \end{aligned} \tag{13}$$

It was shown in [1] that $\Big[I - Q^{-\frac{1}{2}}X(X^\top Q^{-1}X)^{-1}X^\top Q^{-\frac{1}{2}}\Big]$ is idempotent and hence positive semi-definite. Since $T$ is a $p \times p$ diagonal matrix whose $j^{th}$ diagonal element is $(\lambda^2\tau_j^2)^{-1}$, $T \succeq 0$, and hence $(X^\top Q^{-1}X)^{-1} \succeq (X^\top Q^{-1}X + T^{-1})^{-1}$ [2, 3], which implies that

$$0 \le z^\top\Big[I - Q^{-\frac{1}{2}}X(X^\top Q^{-1}X)^{-1}X^\top Q^{-\frac{1}{2}}\Big]z \le z^\top\Big[I - Q^{-\frac{1}{2}}X(X^\top Q^{-1}X + T^{-1})^{-1}X^\top Q^{-\frac{1}{2}}\Big]z \tag{14}$$

for any $z$, showing positive semi-definiteness. To prove positive definiteness, we show that its determinant is non-zero. Let $n = d$, denote $\Lambda$ as the $(d+p)\times(d+p)$ augmented matrix which is written as

$$\begin{aligned} \Lambda^\top\Lambda &= \begin{bmatrix} Y^\top Q^{-\frac{1}{2}} & 0 \\ X^\top Q^{-\frac{1}{2}} & T^{-\frac{1}{2}} \end{bmatrix} \begin{bmatrix} Q^{-\frac{1}{2}}Y & Q^{-\frac{1}{2}}X \\ 0 & T^{-\frac{1}{2}} \end{bmatrix} \\ &= \begin{bmatrix} Y^\top Q^{-1}Y & Y^\top Q^{-1}X \\ X^\top Q^{-1}Y & X^\top Q^{-1}X + T^{-1} \end{bmatrix}. \end{aligned}$$

Note that

$$\begin{aligned} |\Lambda| &= |Q^{-\frac{1}{2}}Y||T^{-\frac{1}{2}} - 0| \\ &= |Q^{-\frac{1}{2}}||Y||T^{-\frac{1}{2}}|, \end{aligned} \tag{15}$$

where $|T^{-\frac{1}{2}}|$ and $|Q^{-\frac{1}{2}}|$ are non-zero. The set of $y$s that lead to linear dependencies among the columns of $Y$ has $\mu^{dn}$ measure zero, implying that $|Y|$ is non-zero $a.e.(\mu^{dn})$. $|\Lambda^\top\Lambda| = |\Lambda^\top||\Lambda| \neq 0$ then gives

$$\begin{aligned} 0 \neq |\Lambda^\top\Lambda| &= |X^\top Q^{-1}X + T^{-1}||Y^\top Q^{-1}Y - Y^\top Q^{-1}X(X^\top Q^{-1}X + T^{-1})^{-1}X^\top Q^{-1}Y| \\ &= |X^\top Q^{-1}X + T^{-1}||Y^\top Q^{-1}Y - \mu^\top\Omega^{-1}\mu| \\ &= |\Omega|^{-1}|Y^\top Q^{-1}Y - \mu^\top\Omega^{-1}\mu|, \end{aligned} \tag{16}$$

which implies that $|Y^\top Q^{-1}Y - \mu^\top\Omega^{-1}\mu| \neq 0.$, and along with positive semi-definiteness, we have $|Y^\top Q^{-1}Y - \mu^\top\Omega^{-1}\mu| > 0 \quad a.e.(\mu^{dn})$. From equation (12) and the inverse Wishart normalizing constant

$$\begin{aligned} &\int_S \int_{\mathbb{R}^{dp}} f(y, q|\beta, \Sigma)\pi(\beta|\Sigma, \tau, \lambda)\pi(\Sigma)\, d\beta^\top d\Sigma \\ &= \frac{|\Omega|^{\frac{d}{2}}\Big[\prod_{i=1}^n h(q_i)\Big]\Big[\prod_{k=1}^d \Gamma(\frac{1}{2}(n+1-k))\Big]}{\pi^{\frac{d(2n-d+1)}{4}}|Q|^{\frac{d}{2}}|T|^{\frac{d}{2}}|Y^\top Q^{-1}Y - \mu^\top\Omega^{-1}\mu|^{\frac{n}{2}}}. \end{aligned}$$

So that

$$\int_{\mathbb{R}_+^{n+p+2}} \int_S \int_{\mathbb{R}^{dp}} f(y,q|\beta,\Sigma)\pi(\beta|\Sigma,\tau,\lambda)\pi(\Sigma)\pi(\tau)\pi(\lambda)\pi(v)\; d\beta^\top d\Sigma dq d\tau d\lambda dv$$

$$= \int_{\mathbb{R}_+^{n+p+2}} \frac{|\Omega|^{\frac{d}{2}} \Big[\prod_{i=1}^n h(q_i)\Big]\Big[\prod_{k=1}^d \Gamma(\frac{1}{2}(n+1-k))\Big]}{\pi^{\frac{d(2n-d+1)}{4}} |Q|^{\frac{d}{2}} |T|^{\frac{d}{2}} |Y^\top Q^{-1} Y - \mu^\top \Omega^{-1}\mu|^{\frac{n}{2}}} \pi(\tau)\pi(\lambda)\pi(v) dq d\tau d\lambda dv. \tag{17}$$

Recalling that the current case is $n = d$ and using (15) and (16), we can rewrite (17) as

$$\int_{\mathbb{R}_+^{n+p+2}} \frac{\Big[\prod_{i=1}^n h(q_i)\Big]\Big[\prod_{k=1}^d \Gamma(\frac{1}{2}(n+1-k))\Big]}{\pi^{\frac{d(d+1)}{4}} |Y|^d} \pi(\tau)\pi(\lambda)\pi(v) dq d\tau d\lambda dv. \tag{18}$$

The expression (18) is proportional to $\Big[\prod_{i=1}^n h(q_i)\Big]$ as a function of $q$ and finite almost surely $\mu^{dn}$, implying that

$$\begin{aligned} c(y) =& \int_{\mathbb{R}_+^{n+p+2}} \int_S \int_{\mathbb{R}^{dp}} f(y,q|\beta,\Sigma)\pi(\beta|\Sigma,\tau,\lambda)\pi(\Sigma)\pi(\tau)\pi(\lambda)\pi(v)\; d\beta^\top d\Sigma dq d\tau d\lambda dv \\ =& \frac{\prod_{k=1}^d \Gamma(\frac{1}{2}(n+1-k))}{\pi^{\frac{d(d+1)}{4}}} < \infty \quad a.e.(\mu^{dn}). \end{aligned}$$

Thus, when $n = d$, the posterior is proper $a.e.(\mu^{dn})$. This also guarantees that the posterior is proper for larger sample sizes; i.e., the posterior is proper for $n > d.$, due to a theorem in [4], which shows when a posterior is proper for a sample size n, it is also proper for all sample sizes larger than $n$. Since the posterior is proper for $n = d$, it also proper when $n > d$ [4].

□

### 2 Derivation of Metropolis-Hastings-within-Gibbs sampler

The original horseshoe prior representation does not yield efficient sampling from the the posterior distribution of the regression coefficients due to non-closed form posterior distributions for the hyper-parameters $(\tau_1^2, \ldots, \tau_p^2)$ and $\lambda^2$. Here we use an alternative sampling scheme [5] based on latent variables that leads to conjugate full conditionals for all parameters and the model becomes easier to solve.

We implement the following scale mixture representation of the half-Cauchy distribution. Let $s$ and $t$ be random variables such that

$$\begin{aligned} s^2|t &\sim \mathcal{IG}(1/2, 1/t), \\ t &\sim \mathcal{IG}(1/2, 1/u^2), \end{aligned} \tag{19}$$

then $s \sim \mathcal{C}^+(0,u)$ [6], where $\mathcal{IG}$ denotes the inverse-gamma distribution, whose pdf is

$$f(x|\alpha,\beta) = \frac{\beta^\alpha}{\gamma(\alpha)} x^{-\alpha-1} \text{exp}\Big(-\frac{\beta}{x}\Big). \tag{20}$$

Our revised hierarchical model using the representation (19) is given by

$$\begin{aligned}
\boldsymbol{Y}_{ij}|q_{ij},\beta,\Sigma &\sim \mathcal{N}(\beta\boldsymbol{X}_i(t_{j-1}),\Sigma/q_{ij}),\\
q_{ij} &\sim \mathcal{G}(v/2,v/2),\\
\pi_1(v) &\propto \Big(\frac{v}{v+3}\Big)^{1/2}\Big\{g'\Big(\frac{v}{2}\Big)-g'\Big(\frac{v+1}{2}\Big)-\frac{2(v+3)}{v(v+1)^2}\Big\}^{1/2},\ \ v>0\\
\beta|\Sigma,\tau^2,\lambda^2 &\sim \prod_{k=1}^{p}\mathcal{N}(0,\lambda^2\tau_k^2\Sigma),\\
\pi_2(\Sigma) &\propto |\Sigma|^{-\frac{d+1}{2}},\\
\tau_k^2|\omega_k &\sim \mathcal{IG}(1/2,1/\omega_k),\ \ k=1,\dots,p\\
\lambda^2|\psi &\sim \mathcal{IG}(1/2,1/\psi),\\
\omega_1,\dots,\omega_p,\psi &\sim \mathcal{IG}(1/2,1).
\end{aligned} \tag{21}$$

#### 2.1 Full conditional for $\beta$

The full conditional distribution for $\beta$ can be obtained as follows

$$\pi(\beta|\cdot)\propto f(y|q,\beta,\Sigma)\pi(\beta|\Sigma,\tau_k^2,\lambda^2)$$

From (10), as a function of $\beta$,

$$\pi(\beta|\cdot)\propto \exp\Big\{-\frac{1}{2}\mathrm{tr}\Big[\Sigma^{-1}(\beta^\top-\mu^\top)\Omega^{-1}(\beta^\top-\mu^\top)^\top\Big]\Big\}. \tag{22}$$

Using the results from (11) regarding the matrix normal density, the full conditional for regression coefficients is given by $\beta|\cdot\sim$ Matrix Normal$(\mu^\top,\Sigma,\Omega)$, where $\mu=(X^\top Q^{-1}X+T^{-1})^{-1}X^\top Q^{-1}Y$ and $\Omega=(X^\top Q^{-1}X+T^{-1})^{-1}$.

#### 2.2 Full conditional for $\Sigma$

The full conditional distribution for $\Sigma$ can be derived as follows

$$\pi(\Sigma|\cdot)\propto f(y|q,\beta,\Sigma)\pi(\beta|\Sigma,\tau_k^2,\lambda^2)\pi(\Sigma).$$

From (12), as a function of $\Sigma$,

$$\pi(\Sigma|\cdot)\propto|\Sigma|^{-\frac{n+d+1}{2}}\exp\Big\{-\frac{1}{2}\mathrm{tr}\Big[\Sigma^{-1}(Y^\top Q^{-1}Y-\mu^\top\Sigma^{-1}\mu\Big]\Big\}. \tag{23}$$

One may apply the results concerning the inverse Wishart density. So, the full conditional for the covariance matrix is distributed according to the inverse Wishart distribution and given by $\Sigma|\cdot\sim \mathrm{IW}(n,\Phi^{-1})$, where $\Phi=(Y^\top Q^{-1}Y-\mu^\top\Omega^{-1}\mu)$.

#### 2.3 Full conditional for $q$

To derive the full conditional for $q$ we use the following representation

$$\pi(q_i|\cdot) \propto f(y_i|q_i,\beta,\Sigma)h(q_i)$$

Thus, as a function of $q_i$,

$$\pi(q_i|\cdot) \propto q_i^{\frac{v+d}{2}} \exp\left\{ -\frac{q_i\left[(\beta^\top x_i - y_i)^\top \Sigma^{-1}(\beta^\top x_i - y_i) + v\right]}{2}\right\} \tag{24}$$

From the density of Gamma distribution, the full conditional for the latent variable $(q_i)$ is given by $q|\cdot \sim \text{Gamma}\Big(\frac{v+d}{2}, \frac{v+(\beta^\top x_i - y_i)^\top\Sigma^{-1}(\beta^\top x_i - y_i)}{2}\Big)$.

#### 2.4 Full conditional for $\tau_k^2$

$$\begin{aligned}
\pi(\tau_k^2|\cdot) &\propto \pi(\tau_k^2|\omega_k)\pi(\beta|\Sigma,\tau_k^2,\lambda^2)\\
&\propto \big(\tau_k^2\big)^{-\frac{1}{2}-1}\exp\left[-\frac{1}{\omega_k\tau_k^2}\right]\big(\tau_k^2\big)^{-\frac{d}{2}}\exp\left[-\frac{1}{2}\big(\tau_k^2\lambda^2\big)^{-1}\beta_k^\top\Sigma^{-1}\beta_k\right]\\
&\propto \big(\tau_k^2\big)^{-\frac{d+1}{2}-1}\exp\left[-\frac{1}{\tau_k^2}\left(\frac{1}{\omega_k}+\frac{1}{2}\big(\lambda^2\big)^{-1}\beta_k^\top\Sigma^{-1}\beta_k\right)\right]
\end{aligned} \tag{25}$$

So, $\tau_k^2|\cdot \sim \mathcal{IG}\Big(\frac{d+1}{2}, \frac{1}{\omega_k}+\frac{1}{2}\big(\lambda^2\big)^{-1}\beta_k^\top\Sigma^{-1}\beta_k\Big)$, $k = 1,\ldots,p$.

#### 2.5 Full conditional for $\lambda^2$

$$\begin{aligned}
\pi(\lambda^2|\cdot) &\propto \pi(\lambda^2|\psi)\pi(\beta|\Sigma,\tau_k^2,\lambda^2)\\
&\propto \big(\lambda^2\big)^{-\frac{1}{2}-1}\exp\left[-\frac{1}{\lambda^2}\right]\prod_{k=1}^{p}\big(\lambda^2\big)^{-\frac{d}{2}}\exp\left[-\frac{1}{2}\big(\tau_k^2\lambda^2\big)^{-1}\beta_k^\top\Sigma^{-1}\beta_k\right]\\
&\propto \big(\lambda^2\big)^{-\frac{pd+1}{2}-1}\exp\left[-\frac{1}{\lambda^2}\left(\frac{1}{\psi}+\frac{1}{2}\sum_{k=1}^{p}\frac{\beta_k^\top\Sigma^{-1}\beta_k}{\tau_k^2}\right)\right]
\end{aligned} \tag{26}$$

Thus, $\lambda^2|\cdot \sim \mathcal{IG}\Big(\frac{pd+1}{2}, \frac{1}{\psi}+\frac{1}{2}\sum_{k=1}^{p}\frac{\beta_k^\top\Sigma^{-1}\beta_k}{\tau_k^2}\Big)$

#### 2.6 Full conditional for $\omega_k$ and $\psi$

From [5], the full conditionals for $\omega_j$ and $\psi$ are given by

$$\begin{aligned}
\omega_k|\cdot &\sim \mathcal{IG}\Big(1, 1+\frac{1}{\tau_k^2}\Big) \quad \text{for} \quad k=1,\ldots,p\\
\psi|\cdot &\sim \mathcal{IG}\Big(1, 1+\frac{1}{\lambda^2}\Big).
\end{aligned}$$

### 3 Analysis of T-cell activation data

Table 1: List of Genes for T-cell activation data

| Gene number | Gene name | Gene number | Gene name |
|---|---|---|---|
| 1 | RB1 | 31 | CDC2 |
| 2 | CCNG1 | 32 | SOD1 |
| 3 | TRAF5 | 33 | CCNA2 |
| 4 | CLU | 34 | PIG3 |
| 5 | MAPK9 | 35 | IRAK1 |
| 6 | SIVA | 36 | SKIIP |
| 7 | CD69 | 37 | MYD88 |
| 8 | ZNFN1A1 | 38 | CASP4 |
| 9 | IL4R | 39 | TCF8 |
| 10 | MAP2K4 | 40 | API2 |
| 11 | JUND | 41 | GATA3 |
| 12 | LCK | 42 | RBL2 |
| 13 | SCYA2 | 43 | C3X1 |
| 14 | RPS6KA1 | 44 | IFNAR1 |
| 15 | ITGAM | 45 | FYB |
| 16 | CTNNB1 | 46 | IL2RG |
| 17 | SMN1 | 47 | CSF2RA |
| 18 | CASP8 | 48 | MPO |
| 19 | E2F4 | 49 | API1 |
| 20 | PCNA | 50 | CYP19 |
| 21 | CCNC | 51 | CIR |
| 22 | PDE4B | 52 | CASP7 |
| 23 | IL16 | 53 | MAP3K8 |
| 24 | APC | 54 | JUNB |
| 25 | ID3 | 55 | IL3RA |
| 26 | SLA | 56 | NFKBIA |
| 27 | CDK4 | 57 | LAT |
| 28 | EGR1 | 58 | AKT1 |
| 29 | TCF12 | | |
| 30 | MCL1 | | |

Table 2: Significant connections among genes for T-cell activation data. Up-regulatory effects are shown in blue and down-regulatory effects in red.

| Gene | Outgoing Connections | Incoming Connections |
|---|---|---|
| 1 | 30 | |
| 7 | -28 | -28 ,-30 |
| 8 | -18 ,28 ,-38 ,-44 ,-50 ,-58 | 11 |
| 11 | 8 | -28 |
| 15 | 27 | 27 |
| 16 | 45 | |
| 17 | -54 | |
| 18 | | -8 |
| 21 | 25 | |
| 22 | | -55 |
| 23 | 52 | -28 ,-30 |
| 25 | | 21 ,46 |
| 26 | | 48 |
| 27 | 15 ,43 ,-45 | 15 |
| 28 | -7 ,-11 ,-23 ,-30 ,-40 ,-54 | -7 ,8 ,29 ,-30 |
| 29 | 28 | -45 ,46 |
| 30 | -7 ,-23 ,-28 ,-33 ,-42 | 1 ,-28 ,48 |
| 32 | 55 | |
| 33 | | -30 |
| 34 | | -55 |
| 38 | | -8 |
| 40 | | -28 ,45 |
| 41 | | -45 |
| 42 | | -30 |
| 43 | | 27 |
| 44 | | -8 |
| 45 | -29 ,40 ,-41 | 16 ,-27 ,46 |
| 46 | 25 ,29 ,45 ,54 | |
| 48 | 26 ,30 | |
| 50 | | -8 |
| 52 | | 23 |
| 53 | | -55 |
| 54 | | -17 ,-28 ,46 |
| 55 | -22 ,-34 ,-53 ,-56 | 32 |
| 56 | | -55 |
| 58 | | -8 |
